# Sex-dependent DNA methylation in amyotrophic lateral sclerosis

**DOI:** 10.64898/2026.09.30.755680

**Authors:** Young Jun Kim, Alfredo Iacoangeli, Ramona A. J. Zwamborn, Wouter van Rheenen, Ross Byrne, Kristel R. Van Eijk, Sarah Marzi, David Watson, Patrick Vourc’h, Philippe Corcia, Mamede de Carvalho, Marc Gotkine, Monica Povedano, Jesus S. Mora Pardina, Fleur Garton, Allan McRae, Pamela J. Shaw, John E. Landers, Jonathan D. Glass, Chris E. Shaw, Nazli A. Başak, Orla Hardiman, Philip Van Damme, Russell L. McLaughlin, Leonard H. van den Berg, Project MinE ALS Sequencing Consortium, Jonathan Mill, Ammar Al-Chalabi, Ahmad Al Khleifat

## Abstract

Amyotrophic lateral sclerosis (ALS) shows well-established sex differences in incidence, age at onset and progression, but their biological basis is unclear. We hypothesised that the DNA methylation changes associated with ALS differ between the sexes, which would point to sex-specific disease processes. To test this, we performed a blood-based epigenome-wide association meta-analysis of ALS cases and controls from Project MinE, an international ALS genomics consortium, and an independent Australian cohort, comprising 8,946 individuals after quality control (5,129 males and 3,817 females; 8,942 with complete covariates in the primary models). We analysed 781,648 autosomal CpGs using complementary sex-stratified case–control models and an ALS-by-sex interaction model that directly tested whether ALS-associated methylation effects differed by sex. Sex-stratified analyses identified no genome-wide significant differences between male and female ALS effects. In contrast, interaction testing identified seven robust sex-dependent ALS-associated differentially methylated positions (*P* < 9 × 10^−8^), mapping to *KCNQ1, ADPRH, HLA-F, GHRLOS, IRF5, RAD51B* and one intergenic region. The strongest signal was at *KCNQ1* (*P* = 3.1 × 10^−9^). Six of the seven remained genome-wide significant in a four-stratum sensitivity analysis, and none was detected in previous conventional ALS or population sex-associated methylation analyses. Together, these findings suggest a distinct sex-conditional component of the ALS blood methylome, implicating immune, metabolic and ion-channel loci and providing new candidates for investigating biological heterogeneity between males and females with ALS.

## Introduction

Amyotrophic lateral sclerosis (ALS) is a neurodegenerative disease characterised by the progressive loss of upper and lower motor neurons, leading to muscle weakness, paralysis and, ultimately, respiratory failure [1, 2]. A family history of ALS or frontotemporal dementia is obtained in about 10% of people with ALS (reported range 5–20%), and a monogenic cause has been identified in about 80% of these [3]. Monogenic ALS is not confined to familial disease: about 15–20% of people with apparently sporadic ALS also carry a Mendelian ALS gene variant [3], so most people with a monogenic cause of ALS have no family history. Moreover, many genes implicated in familial ALS have also been associated with apparently sporadic disease [4], and rare risk variants are found in more than 20% of all people with ALS [5]. Beyond these large-effect variants, twin studies estimate the heritability of ALS at around 60%, largely reflecting additive, polygenic risk [6]. These observations have motivated gene–environment and multistep exposome models, in which lifetime environmental exposures act on this genetic background during ageing to precipitate disease [7, 8, 9].

Epigenetic mechanisms provide a plausible interface between the environment and gene regulation. DNA methylation (DNAm), the addition of a methyl group to cytosine, predominantly at CpG dinucleotides, is the most widely studied epigenetic mark and can both regulate gene activity and record past exposures. Blood-based epigenome-wide association studies (EWAS) therefore offer a scalable route to interrogate the molecular correlates of ALS. As a durable record of such exposures, DNAm offers a tractable entry point into the non-genetic architecture of non-monogenic ALS; yet the epigenetic layer arising from gene–environment interaction, and its stratification by host factors, has been comparatively overlooked [10]. Across blood and central nervous system tissues, ALS is associated with a global increase in 5-methylcytosine content and with altered methylation of genes enriched for immune, inflammatory and metabolic pathways [11]. The largest such study to date was a case–control EWAS meta-analysis in 9,706 samples and identified differentially methylated positions enriched for metabolic, cholesterol-biosynthesis and inflammatory pathways, several of which were also associated with survival [12]. This established both a robust methodological framework and a reference set of ALS-associated methylation signals in whole blood. Since then, ALS methylation studies have scaled to multi-cohort, cross-population and multi-omics designs, and have broadened beyond mean methylation to features such as epigenetic drift, rare epivariations and survival association [13, 14].

ALS displays well-documented sexual dimorphism in incidence, age at onset and progression rate [1]: men are affected more frequently than women (male-to-female ratio ≈1.3–1.5:1) and tend to develop disease at a younger age. These differences have been linked to sex-steroid signalling as well as to sex-specific genetic and environmental factors: steroid hormone receptors are expressed in motor neurons and glia and become dys-regulated with disease progression in ALS models [15], and Mendelian-randomisation analyses suggest that higher bioavailable testosterone is associated with lower ALS risk [16]. DNAm is itself strongly sex-dependent; while sex differences on the X and Y chromosomes are expected (X-inactivation, Y-linked probes), autosomal sex-associated differentially methylated positions (saDMPs) are of greater biological interest, and have been catalogued in the general population, in whole blood [17] and in plasma cell-free DNA [18]. Although sex differences in DNAm have been documented in other neurodegenerative disorders, they remain under-investigated in ALS: most ALS EWAS match on or statistically adjust for sex rather than test it explicitly, and few report sex-stratified comparisons [11]. Where this has been examined in other brain disorders, sex-stratified EWAS have improved discovery over sex-adjusted models and revealed a sex-by-disease interaction on methylation, arguing for modelling sex explicitly rather than adjusting it away [19]. Whether, and how, the ALS methylation signal itself differs between the sexes therefore remains an open question.

Here we test the hypothesis that the DNA methylation changes associated with ALS differ between the sexes, and that such differences are detectable in peripheral blood. We conduct a large, blood-based, sex-associated EWAS of ALS in which sex is treated not as a nuisance covariate to be adjusted away but as a biological axis to be resolved within the ALS methylome. Using blood DNAm from Project MinE, an international ALS genomics consortium [20], profiled on the 450k and EPIC arrays, and from an independent Australian cohort, and applying the quality-control and association pipeline of our previous work [12], we directly compare two modelling strategies for sex-specific effects: (i) a sex-stratified case–control EWAS with a formal test for a difference in the ALS effect between sexes, and (ii) a sex-by-case–control interaction model. By isolating the sex-specific component that adjustment-based analyses discard, this design seeks a stratifiable slice of the ALS methylome relevant to the disease’s sex bias in incidence, onset and progression.

## Materials and Methods

### Study population and cohorts

We analysed blood-derived DNA methylation from participants with ALS (diagnosed with definite, probable, or probable laboratory-supported ALS according to the revised El Escorial criteria [21]) and population-based controls from Project MinE, profiled on the Illumina 450k and EPIC arrays, and an independent Australian cohort from the Strategic ALS Australia–Systems Genomics Consortium (SALSA-SGC) [22], profiled on the Illumina EPIC array. No participants were excluded on the basis of family history or ALS gene-variant status in this analysis, and *C9orf72* repeat-expansion carriers were retained (Table 1); recruitment criteria nevertheless differed between contributing cohorts, some of which excluded people with a family history of ALS while others included or specifically recruited familial cases [12]. After quality control, the primary three-stratum analysis comprised 8,946 individuals (3,817 females and 5,129 males; 6,287 ALS cases and 2,659 controls) across three strata: Project MinE 450k, Project MinE EPIC and the Australian AUS1 stratum. The Australian data form a single dataset split at the median red/green signal-intensity ratio (threshold 0.65) into AUS1 (*n* = 1,087) and a smaller AUS2 stratum (*n* = 168), following our previous work [12], with an intensity-appropriate normalisation applied to each (AUS1, `nanes`; AUS2, `dasen`). Because AUS2 became very sparse once partitioned by sex and case/control status, it was excluded from the primary analysis and retained only in a four-stratum sensitivity analysis (*n* = 9,114). Per-stratum counts are given in Table 1. Participants were predominantly of European ancestry. All contributing studies were conducted with the approval of the relevant institutional review boards or research ethics committees, and all participants gave written informed consent, in accordance with the Declaration of Helsinki.

**Table 1:** Study population after quality control, by stratum, sex and case/control status. The primary analysis uses three strata (MinE 450k, MinE EPIC and AUS1; *n* = 8,946); AUS2 is included only in the four-stratum sensitivity analysis. Age is DNAm-predicted age (*pred_age*) in years, mean (SD), the age covariate used in the models; real age at blood draw is available for the Project MinE strata, whereas AUS1 provides only predicted age, which is markedly more variable and should be read with caution. Site of onset (spinal / bulbar) and *C9orf72* repeat-expansion carriers are counted among cases and shown for the Project MinE strata; *C9orf72* status was unavailable for 413 MinE EPIC cases, and neither field was available for AUS1.

|  | MinE 450k | MinE EPIC | AUS1 | AUS2 | Total |
| --- | --- | --- | --- | --- | --- |
| Female, control | 629 | 406 | 267 | 9 | 1,311 |
| Female, case | 1,184 | 1,112 | 219 | 66 | 2,581 |
| Male, control | 781 | 350 | 226 | 11 | 1,368 |
| Male, case | 1,760 | 1,637 | 375 | 82 | 3,854 |
| Stratum total | 4,354 | 3,505 | 1,087 | 168 | 9,114 |
| Mean age, years | 62.6 (9.7) | 63.0 (8.5) | 66.2 (25.0) | – | 63.2 (12.3) |
| Onset spinal / bulbar | 1,899 / 844 | 1,862 / 683 | – | – | 3,761 / 1,527 |
| *C9orf72* expanded (cases) | 183 | 123 | – | – | 306 |

### Array processing and quality control

Analyses started from the quality-controlled Project MinE *β*/phenotype matrices, generated with *minfi* [23] and *wateRmelon* [24] following the Project MinE methylation pipeline of our previous work [12]. Samples were excluded on median methylated/unmethylated intensities, reported-versus-predicted sex discordance, bisulfite conversion efficiency (bscon), control-probe metrics (*MethylAid*) [25], detection-*P* /bead-count metrics and control-probe PCA outliers; signal intensities were normalised with `dasen`, except the dye-biased Australian AUS1 stratum, which was normalised with `nanes` [24]; and cross-reactive and SNP-affected probes were removed. Each stratum was then restricted to autosomal CpGs (chromosomes 1–22); probes missing in more than 5% of that stratum’s samples were dropped, and the small residual fraction of missing values (≈ 0.1–0.2%) was imputed by per-CpG (row) means, leaving 781,648 autosomal CpGs for analysis. Because cross-reactive and common-SNP-affected probes had already been removed, the reported associations are unlikely to reflect probe cross-hybridisation or genotype at the interrogated site; a formal methylation-quantitative-trait-locus analysis to exclude distal genetic effects was left to future work. *β*-values were converted to M-values by the logit transform after clamping to [10^*−*6^, 1 − 10^*−*6^].

### Cell-composition estimation

White-blood-cell proportions were estimated by the reference-based Houseman method [26]. Of the six estimated cell types, B cells were dropped to avoid compositional collinearity and the remaining five (CD4^+^ T cells, CD8^+^ T cells, natural killer cells, monocytes and granulocytes) were included as covariates, matching the covariate set of our previous work [12].

### Statistical models

Methylation was analysed at each autosomal CpG using *β*-values as the primary response, for interpretability on the methylation scale, with M-values (logit-transformed *β*) analysed as a sensitivity; the per-CpG response is denoted *y* below. Autosomes were the primary analysis; the sex chromosomes were analysed and reported separately, because X-inactivation and Y-linked probes dominate any genome-wide sex contrast. Both models, and the designation of sex as the effect modifier of interest, were specified a priori; the two are complementary. Both were fitted on complete covariate cases: any sample with a missing value in a model covariate (predicted age, batch, the estimated cell proportions or the residual principal components) was excluded before fitting, because the design matrix would otherwise drop it silently and break alignment with the methylation matrix. This affected only the Australian strata and is reconciled against the quality-control counts in the Results.

#### Model 1: sex-stratified EWAS

Within each stratum and separately within each sex, we fitted an ordinary least-squares case/control EWAS per CpG (*y* ∼ ALS + covariates; *limma* `lmFit` without empirical-Bayes moderation) [27]. Covariates were predicted (DNAm) age, technical batch, the five estimated cell proportions and a tuned number of array-wide residual principal components; for the two Project MinE strata a further 30 control-probe principal components were included (these are unavailable for the Australian strata, which are supplied as *β*-matrices without control-probe intensities). Per-stratum statistics were corrected with *bacon* [28] and combined, within each sex, across strata by inverse-variance fixed-effect meta-analysis [29]. A formal *z*-test for a between-sex difference in the ALS effect (*β*_male_ = *β*_female_) was then computed per CpG; positions with *p*_diff_ *<* 9 × 10^*−*8^ were called sex-associated DMPs (saDMPs). Because the ALS effect is estimated separately within each sex, this model also yields the per-sex direction of change (hyper- or hypo-methylation) at each saDMP.

#### Model 2: ALS-by-sex interaction

In the full sample (both sexes, cases and controls) we fitted, per CpG, *y* ∼ ALS × Sex_*M*_ + covariates, retaining both main effects for marginality and using the same per-stratum covariate set as Model 1. The `ALS:Sex_M` interaction coefficient, the difference in the ALS methylation effect between males and females, is the headline test. Per-stratum interaction statistics were bacon-corrected and combined by inverse-variance fixed-effect meta-analysis, with between-stratum heterogeneity assessed by Cochran’s *Q*; positions with *P <* 9 × 10^*−*8^ were called saDMPs. Note that the sign of this interaction alone does not indicate which sex is hyper- or hypo-methylated; the per-sex direction is resolved from the sex-stratified estimates (Model 1).

#### Inflation control and PC tuning

Following our previous work [12], the number of array-wide residual principal components was tuned per fit over the grid {5, 10, 15, 20, 25, 30} with a floor of 10, selecting the smallest number whose pre-*bacon* inflation factor met the target, and falling back to the largest value in the grid where no value reached it, as happened for AUS1 in both models; *bacon* [28] was then applied to the final fit (`set.seed`(10)). For the sex-stratified case–control fits (Model 1) the target was the sample-size-normalised *λ*_1000_ ≤ 1.15; for the interaction fits (Model 2), for which *λ*_1000_ is undefined, the conventional *λ* ≤ 1.10 was used. *λ* is reported before and after *bacon* correction.

#### Direction and magnitude of effects

For each sex-associated DMP the direction of change, that is whether ALS is associated with a gain (hyper-methylation) or loss (hypo-methylation) and in which sex, was read from the sign of the sex-stratified ALS estimates (*β*_male_ and *β*_female_ from Model 1), because the interaction coefficient alone is sign-ambiguous with respect to sex. As the primary analysis is on the *β* scale, these estimates are Δ*β* values (the case/control difference in mean *β*) and are reported directly. The M-value analysis, which is robust to the heteroscedasticity of *β*-values [30], served as a sensitivity check and was sign-concordant with the primary analysis at all seven saDMPs.

### Meta-analysis across strata

Per-CpG effect sizes and standard errors were combined by inverse-variance fixed-effect meta-analysis [29] (NA-safe alignment, so probes present on only a subset of strata are retained where estimable). A fixed-effect model was chosen a priori because the strata estimate the same biological effect under a single harmonised pipeline; between-stratum heterogeneity was quantified by Cochran’s *Q* and *I*^2^, and a random-effects (DerSimonian– Laird) meta-analysis was run as a sensitivity check. The primary analysis used three strata (MinE 450k, MinE EPIC and AUS1). AUS2 was excluded because, once partitioned by sex and case/control status, its cells become very small (for example 9 female and 11 male controls; Table 1), which inflates the variance of its per-stratum estimates; a four-stratum analysis including AUS2 was retained as a sensitivity check.

### Significance, annotation and software

Genome-wide significance was set at *P <* 9 × 10^*−*8^ [31], and Benjamini–Hochberg false-discovery-rate values were additionally computed for all tests [32]. saDMPs were annotated to the nearest gene and to CpG island, shore, shelf and open-sea contexts using the Illumina 450k and EPIC manifests (hg19), and characterised for genomic context relative to the array background. All analyses were performed in R 4.5 using *limma* [27], *bacon* [28], *minfi* [23] and *metafor* [29], with *data*.*table* for input/output; *wateRmelon* [24] was used in the upstream QC and normalisation.

## Results

### Cohorts and samples

The primary three-stratum analysis comprised 8,946 individuals (3,817 females and 5,129 males; 6,287 ALS cases and 2,659 controls) across Project MinE 450k, Project MinE EPIC and AUS1; the four-stratum sensitivity analysis additionally included AUS2 (168 samples), giving 9,114 in total (Table 1); reported age is DNAm-predicted age (*pred_age*), the age covariate used throughout. Because analysis began from quality-controlled matrices, per-step exclusion counts were not available to us; the QC procedure and the retained totals (781,648 autosomal CpGs; 8,946 individuals) are given in the Methods. The counts in Table 1 are those passing quality control. Because every model was fitted on complete covariate cases (Methods), a small number of samples with a missing covariate value were not carried into the fitted models: four in AUS1 (three female and one male control) and three in AUS2 (one female and two male cases), with no losses in either Project MinE stratum. In AUS1 these samples lacked the array-wide residual principal components used for inflation control, and most also lacked a DNAm-predicted age. The fitted models therefore comprised 8,942 individuals in the primary three-stratum analysis and 9,107 in the four-stratum sensitivity analysis, which are the sample sizes reported in Table S2.

### Sex-stratified analysis yields no genome-wide saDMPs (Model 1)

The sex-stratified case/control EWAS, followed by a formal test for a difference in the ALS effect between the sexes, did not identify any genome-wide significant saDMPs; the strongest between-sex difference genome-wide reached *p*_diff_ = 1.2 × 10^*−*6^ (cg03270395), and the strongest among the positions identified by the interaction model reached *p*_diff_ ≈ 3 × 10^*−*6^ (at *ADPRH* and *KCNQ1*), short of the 9 × 10^*−*8^ threshold. Inflation was well controlled in the Project MinE strata after PC tuning and *bacon* (post-*bacon λ*_1000_ between 1.02 and 1.03 across sexes), whereas the small AUS1 stratum retained higher residual inflation, most markedly in females (post-*bacon λ*_1000_ = 1.20; 219 cases and 264 controls) and less so in males (1.08; Table S2). At the meta-analysis level each per-sex EWAS was well calibrated (males *λ* = 1.04, females *λ* = 1.06), and the sex-difference test itself showed no residual inflation (*λ* = 1.00; Figure S1A–C).

### An interaction model identifies seven robust saDMPs (Model 2)

In contrast, the ALS-by-sex interaction model, applied to 781,648 autosomal CpGs, identified eight positions at genome-wide significance (*P <* 9 × 10^*−*8^). One of these, cg17051239 (*CCDC121*), was not corroborated by the sex-stratified analysis (Table 2) and was excluded as a likely false positive, leaving seven robust saDMPs (Table 2; Figure 1). Six of the seven were also genome-wide significant in the four-stratum sensitivity analysis that additionally included AUS2, with identical per-sex direction; only cg11587584 (*HLA-F*) was specific to the primary three-stratum analysis. Between-stratum heterogeneity was limited (Cochran’s *Q*): only cg10151367 (*KCNQ1*) reached the Bonferroni-corrected threshold, and then marginally (*p*_het_ = 0.0071 against a threshold of 0.0071; *I*^2^ = 80%), with cg14992144 (*GHRLOS*) heterogeneous at nominal significance only (*p*_het_ = 0.017, *I*^2^ = 75%); the remaining five positions showed no heterogeneity (*p*_het_ ≥ 0.14, *I*^2^ ≤ 49%). After PC tuning (10, 10 and 30 residual principal components for MinE 450k, MinE EPIC and AUS1 respectively) and *bacon* correction, per-stratum inflation in the primary (*β*-scale) analysis fell from pre-*bacon λ* of 1.06, 1.07 and 1.44 to post-*bacon* 1.02, 1.02 and 1.03 (Table S2); the interaction meta-analysis itself was well calibrated (*λ* = 1.05; Figure S1D).

**Table 2:** The seven robust ALS sex-associated DMPs from the primary three-stratum interaction model, ordered by meta-analysis *P* (hg19 coordinates). Δ*β*_int_ is the interaction coefficient (*β* scale) with standard error SE; FDR is the Benjamini–Hochberg false-discovery rate across all tested CpGs; Male and Female Δ*β* are the sex-stratified ALS case/control effects (Model 1); the pattern gives the per-sex direction (hyper/hypo at nominal *p <* 0.05). All are robust to AUS2 inclusion except cg11587584 (*HLA-F*). An eighth position, cg17051239 (*CCDC121*), reached interaction significance but was not corroborated by the between-sex test (*p*_diff_ = 0.96) and is excluded here as a likely false positive. Gene symbols follow current HGNC nomenclature; *RAD51B* is listed under its previous symbol, *RAD51L1*, in the Illumina manifest.

| CpG | Position | Gene | $\Delta\beta_{\text{int}}$ | SE | Meta $P$ | FDR | Male $\Delta\beta$ | Female $\Delta\beta$ | Pattern |
| --- | --- | --- | --- | --- | --- | --- | --- | --- | --- |
| cg10151367 | 11:2847642 | *KCNQ1* | −0.0066 | 0.0011 | $3.1 \times 10^{-9}$ | 0.0014 | −0.0022 | +0.0033 | M hypo / F hyper |
| cg04678315 | 12:125212010 | intergenic | −0.0068 | 0.0012 | $3.9 \times 10^{-8}$ | 0.0061 | −0.0020 | +0.0012 | M hypo / F n.s. |
| cg16904330 | 3:119299162 | *ADPRH* | −0.0060 | 0.0011 | $4.3 \times 10^{-8}$ | 0.0061 | −0.0030 | +0.0023 | M hypo / F hyper |
| cg11587584 | 6:29692372 | *HLA-F* | +0.0074 | 0.0014 | $4.5 \times 10^{-8}$ | 0.0061 | +0.0024 | −0.0040 | M hyper / F hypo |
| cg14992144 | 3:10334743 | *GHRLOS* | −0.0061 | 0.0011 | $4.7 \times 10^{-8}$ | 0.0061 | −0.0008 | +0.0021 | M n.s. / F hyper |
| cg04086097 | 7:128580792 | *IRF5* | −0.0063 | 0.0012 | $5.9 \times 10^{-8}$ | 0.0065 | −0.0007 | +0.0019 | M n.s. / F hyper |
| cg05315321 | 14:68844276 | *RAD51B* | −0.0050 | 0.0009 | $7.1 \times 10^{-8}$ | 0.0069 | −0.0026 | +0.0006 | M hypo / F n.s. |

**Figure 1:**
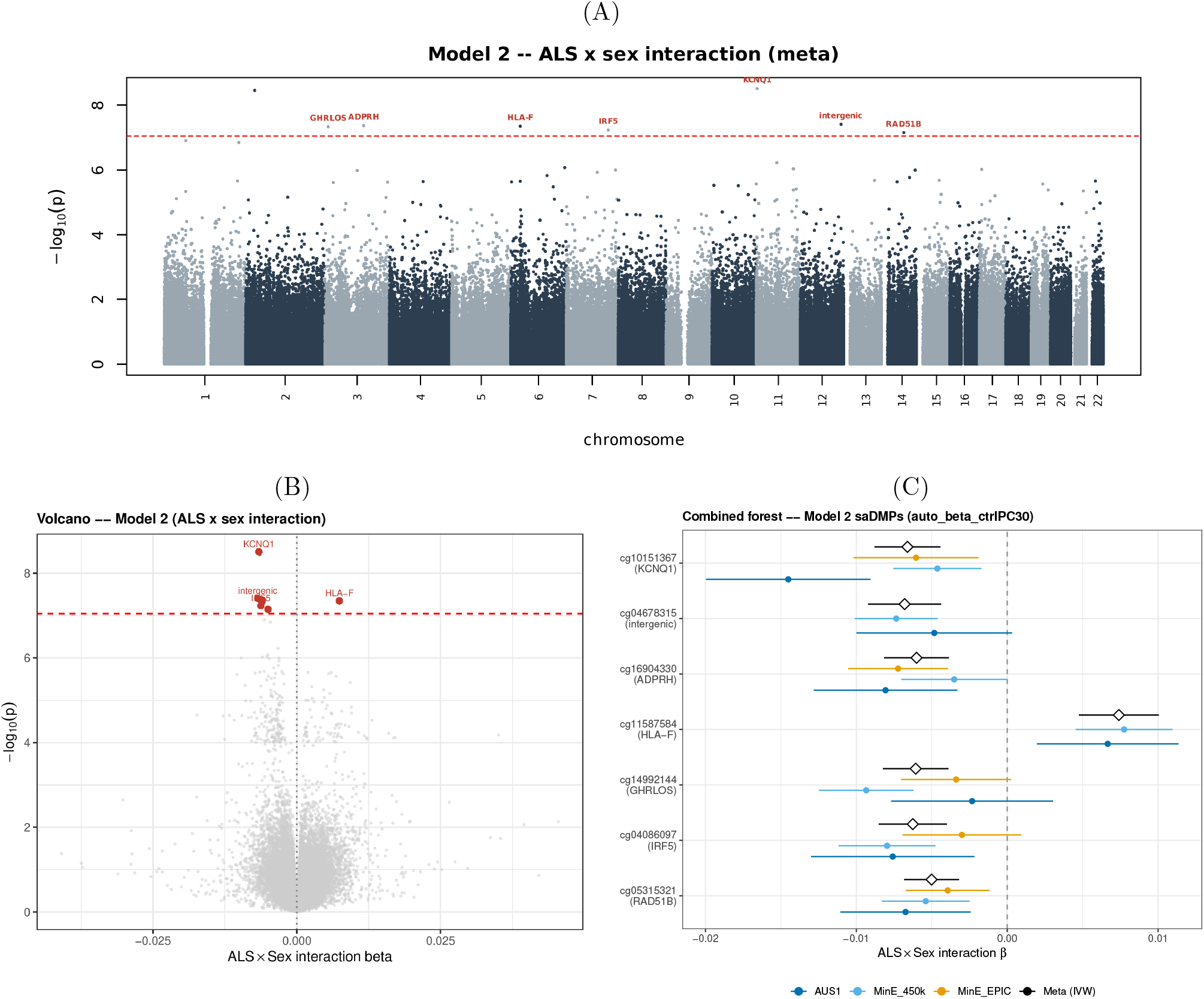
Sex-associated DNA methylation in ALS from the interaction model (Model 2; primary three-stratum analysis, *β* scale). (A) Manhattan plot of the ALS-by-sex interaction meta-analysis across 781,648 autosomal CpGs; the dashed line marks genome-wide significance (*P <* 9×10^*−*8^) and the seven robust saDMPs are labelled by nearest gene. (B) Volcano plot of the interaction effect size against − log_10_ *P* , with the seven saDMPs highlighted. (C) Forest plot of the seven saDMPs showing the per-stratum interaction estimate (MinE 450k, MinE EPIC, AUS1) and the inverse-variance meta-analysis (diamond) with 95% confidence intervals. The excluded position cg17051239 (*CCDC121*) is not labelled.

Decomposing each interaction into its sex-stratified components (Model 1) resolved the direction of effect (Table 2). At most saDMPs the ALS-associated change was in opposite directions in the two sexes, or confined to one sex. At four positions, including *KCNQ1* (cg10151367) and *ADPRH* (cg16904330), methylation was increased in female cases (hyper-methylation) while decreasing or unchanged in males; at *HLA-F* (cg11587584) the pattern was reversed, with hyper-methylation in males and hypo-methylation in females. The excluded position, cg17051239 (*CCDC121*), reached interaction significance although neither per-sex effect was individually significant and the two sexes did not differ on the formal between-sex test (*p*_diff_ = 0.96; per-sex Δ*β* = −0.0002 in both); its interaction signal is therefore not corroborated by the stratified analysis, and it was excluded on that basis. Among the seven robust saDMPs, effect sizes were uniformly small (per-sex |Δ*β*| from 0.0006 to 0.0033, i.e. below 0.4% methylation), with female hyper-methylation the predominant pattern (Figure 2). The seven robust saDMPs mapped to six genes and one intergenic position (cg04678315): *KCNQ1, ADPRH, HLA-F, GHRLOS*, the immune regulator *IRF5* and *RAD51B*.

**Figure 2:**
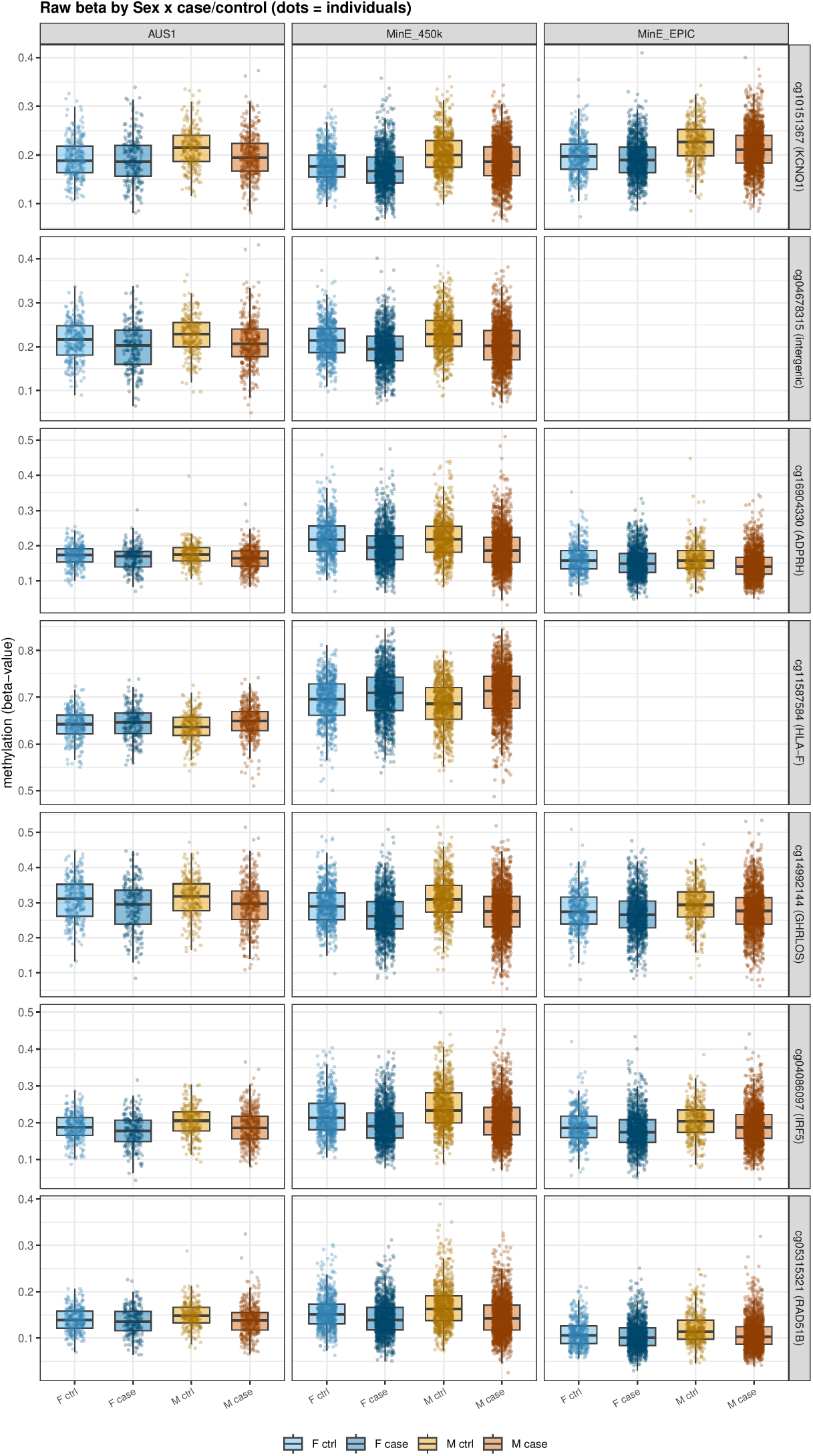
Raw methylation (*β*-value) at the seven robust saDMPs, split by sex and case/control status within each array stratum (dots are individuals). The panels show the per-sex direction of the ALS-associated change that the interaction model captures: female-specific hyper-methylation (*GHRLOS, IRF5*), male-specific hypo-methylation (*RAD51B* and the intergenic probe), and sex-divergent changes (*KCNQ1, ADPRH, HLA-F*).

Effect directions were fully concordant across strata (sign concordance of one for all seven positions, with

95% confidence intervals excluding zero), every stratum contributing an effect of the same sign; magnitudes nonetheless varied, most visibly at *KCNQ1*, where the AUS1 estimate was about twice the meta-analytic one. In leave-one-out meta-analyses none of the seven retained genome-wide significance in every subset (worst-case *P* ranging from 3 × 10^*−*5^ to 0.07); the intergenic probe cg04678315, which was present on only two of the three strata, was the least robust (worst-case *P* = 0.07), followed by *GHRLOS* (cg14992144, worst-case *P* = 0.05). A random-effects meta-analysis gave concordant point estimates. The meta-analytic standard errors were of order 0.001 (Table 2), placing each estimate five to six standard errors from the null, and all seven were sign-concordant on the M-value scale.

### Stratified versus interaction modelling

The interaction test (Model 2) returned genome-wide significant positions where the sex-stratified difference test (Model 1) returned none, the latter reaching *p*_diff_ = 1.2 × 10^*−*6^ at its strongest (Figure S2). None of the seven saDMPs coincided with published general-population sex-associated DMPs [17], either at the level of individual CpGs or of their annotated genes, and none was a genome-wide significant ALS-DMP in our previous blood ALS EWAS meta-analysis [12] (Table S1).

### Genomic characterisation

Relative to the tested background, the saDMPs were not significantly enriched for any CpG-island context or gene region (Fisher’s exact test). They were over-represented at enhancers: two of the seven lay in enhancer regions against 0.26 expected (odds ratio 10.3, Fisher *P* = 0.026). The background for this test was the full set of analysed probes.

### The X chromosome

We repeated both models on the X chromosome (17,594 CpGs). Model 1, which compares cases and controls within each sex, identified no X-linked saDMP (no CpG at FDR *<* 0.05; strongest *p*_diff_ = 6 × 10^*−*5^). The Model 2 interaction showed test-statistic inflation on X (*λ*_GC_ = 1.20, against 1.05 on the autosomes), with a correspondingly large number of nominal hits. Male hemizygosity and X-inactivation give males and females markedly different methylation variance on X, which the common residual variance assumed by the interaction model does not accommodate. All saDMPs reported in this study are autosomal.

## Discussion

This study set out to determine whether the DNA methylation changes associated with ALS differ between the sexes and, if so, whether such differences are detectable in peripheral blood. The principal result is as much methodological as biological. A direct case-by-sex interaction test (Model 2) identified eight autosomal positions at genome-wide significance, of which seven were corroborated by the sex-stratified analysis and retained as robust sex-associated differentially methylated positions, whereas the sex-stratified difference test (Model 1), which compares male and female case-control estimates after the fact, returned none and reached only *p*_diff_ = 1.2 × 10^*−*6^ at its strongest. The two designs ask subtly different questions. The stratified test asks whether the male and female case-control effects differ and pays a heavy power penalty because it compounds the sampling error of two separately estimated effects, whereas the interaction model estimates that difference directly. Interaction terms require substantially larger samples than main effects for equivalent power, because the interaction contrast carries the variance of both estimates, so the small number of significant positions and the null stratified result are expected rather than contradictory. The practical implication is that any sex-specific methylation signal in ALS is modest in magnitude, and that detecting it depends on analytical design as much as on sample size. That none of the seven positions coincided with published general-population sex-associated DMPs [17], at either CpG or gene level, further argues that they are disease-context interactions rather than constitutive sex differences that happen to be present in cases.

Why these positions reflect ALS rather than sex alone is worth spelling out, since the two interpretations are easily conflated. The distinction is built into the interaction test: a methylation difference that is present equally in cases and controls loads onto the sex main effect and cannot, by construction, generate a significant interaction, so a saDMP necessarily marks an ALS case-control effect whose size or direction depends on sex rather than a constitutive sex difference. The sex-stratified decomposition bears this out, as each robust saDMP carries a genuine ALS case-versus-control effect in at least one sex (Table 2), and none is a genome-wide ALS-DMP in a conventional main-effect analysis (Table S1). What we report is therefore the sex-conditional component of the ALS methylome: a signal anchored to disease status within each sex, yet invisible both to a main-effect ALS EWAS and to a general-population sex-DMP scan.

### Biological interpretation of the sex-associated loci

The seven robust positions annotate to six genes and one intergenic region. Nearest-gene assignment is not evidence of regulation, but several of the implicated loci sit within pathways already tied to ALS and, in some cases, to sex-biased physiology. Decomposed into their sex-stratified components, the credible loci fall into three directional patterns: a female-specific change, in which methylation shifts in female cases only (*GHRLOS* and *IRF5*, both hyper-methylated in women); a male-specific change confined to male cases (*RAD51B* and the intergenic probe cg04678315, both hypo-methylated in men); and a sex-divergent pattern in which the sexes move in opposite directions (*KCNQ1* and *ADPRH*, hypo in men and hyper in women, and *HLA-F*, the reverse). We describe direction throughout as the case-versus-control change within each sex, since that is what a sex interaction encodes rather than a single overall effect (Figure S3).

*KCNQ1* (cg10151367) belongs to the Kv7 family of voltage-gated potassium channels that generate the M-current governing neuronal firing. Motor-neuron hyperexcitability is an early and measurable feature of ALS: pharmacological potentiation of Kv7 channels lowers cortical and spinal excitability in patients [33] and is neuroprotective against excitotoxicity in motoneuron models [34]. *KCNQ1* is imprinted and is not itself the neuronal Kv7 paralogue, so the specific relevance of this locus is unproven, but it flags a channel family central to ALS pathophysiology. *GHRLOS* (cg14992144) is the antisense transcript of the ghrelin (*GHRL*) locus. Ghrelin is an orexigenic and neuroprotective hormone, and loss of ghrelin action tracks the weight loss, hypermetabolism and shorter survival characteristic of ALS [35, 36]; because body composition and energy metabolism differ markedly between men and women, a sex-differential signal here is biologically coherent.

Two further positions implicate immune regulation. *IRF5* (cg04086097) is a master transcription factor driving pro-inflammatory macrophage polarisation [37]; *IRF5* is moreover a sex-dimorphic immune locus, showing a gene-by-sex interaction in systemic lupus erythematosus in which affected men carry a higher burden of risk alleles than women [38]. Here methylation increased in female but not male cases; our data do not establish why the effect falls in that direction, and the genetic sex-dimorphism reported at this locus runs the other way, with affected men carrying the greater risk-allele burden. *HLA-F* (cg11587584) is a non-classical MHC class I gene within the HLA region, a locus that shows immune-mediated genetic enrichment in frontotemporal dementia [39]. This locus is notable because a recent sex-stratified ALS genome-wide association study found the HLA locus, which harbours *HLA-F*, to be associated with ALS risk specifically in males [40], which is consistent in direction with the male-biased effect we observe here (hyper-methylation in male cases, hypo-methylation in females), although that association is to the HLA region rather than to this CpG. Neuroinflammation is an established contributor to ALS, and sexual dimorphism in immune function is well documented [41], making these plausible substrates for sex-differential disease effects.

*RAD51B* (*RAD51B*; cg05315321) is a paralogue of the homologous-recombination machinery; impaired DNA-damage response and genomic instability are increasingly viewed as early triggers in ALS [42]. Two of the remaining positions, cg16904330 (*ADPRH*) and the intergenic probe cg04678315, are statistically well supported, with the sexes differing on the stratified test, but have no established connection to ALS and should be regarded as hypothesis-generating. The eighth, cg17051239 (*CCDC121*), is significant in the interaction model yet shows no difference between the sexes on the stratified test (*p*_diff_ = 0.96, with per-sex effects of −0.0002 in both), so its signal is uncorroborated and most plausibly a false positive arising from residual heteroscedasticity rather than a genuine sex-differential effect. That the other seven loci each show a between-sex difference on the stratified test (all *p*_diff_ *<* 0.05) whereas cg17051239 does not suggests that the interaction test is not producing these signals spuriously. Across all seven the absolute effects are small, below one per cent methylation, and rest on the joint meta-analysis rather than any single cohort. Effects of this size sit below the single-probe technical noise of these arrays, which is of order Δ*β* = 0.05; what makes them estimable is not per-probe precision but the meta-analytic standard error over nearly nine thousand samples, of order 0.001, which places each estimate five to six standard errors from the null. That argument justifies reporting the positions, not treating them as established: these loci nominate candidate pathways rather than mechanisms, and require replication and functional validation before mechanistic weight is placed on them.

### What these loci do not explain

Because this is a case-control comparison, it characterises how the epigenetic signature of ALS differs between men and women; it does not address why ALS is more common in men (a male-to-female incidence ratio of roughly 1.3 to 1.5 to one) [1], which is a question of incidence and risk rather than of the disease signature, and the direction of a methylation change does not by itself reveal whether that change is protective or pathogenic. The axes implicated here, immune regulation through *IRF5* and *HLA-F* and energy metabolism through *GHRLOS*, are independently known to be sexually dimorphic and relevant to ALS, but whether any of them contributes to the sex bias in disease susceptibility is beyond what the present design can establish, and would require risk-based, prospective and causal studies. Sex hormones are one plausible upstream mediator of such effects: sex-steroid receptors are expressed in motor neurons and glia and become dysregulated over disease progression in ALS mice [15], and Mendelian-randomisation analyses link sex-hormone exposure to neurodegenerative-disease risk [16]; whether these pathways shape the methylation differences reported here is untested.

### Limitations and future directions

Several limitations qualify these findings and frame the work that should follow. First, methylation was measured in whole blood rather than in the central nervous system. DNA methylation is tissue-specific, so blood-based saDMPs need not reflect the state of motor neurons or glia; blood remains informative as an accessible biomarker and as a readout of the peripheral immune involvement that characterises ALS, but confirmation in central nervous system tissue, or in cell-sorted and disease-proximal samples, is needed before neural interpretation. Second, the effect sizes are small and the design is cross-sectional, so we cannot separate causal epigenetic differences from consequences of the disease or its correlates, including medication, physical decline, smoking and the weight loss noted above; longitudinal sampling before symptom onset and causal-inference approaches such as two-step Mendelian randomisation would help establish direction. Third, the sex-interaction saDMPs were not tested in an independent replication cohort, and nearest-gene annotation is not proof of regulation; independent, sex-balanced ALS cohorts, together with methylation-quantitative-trait and expression mapping and targeted functional assays, are required to link each CpG to a gene and a mechanism. Fourth, the contributing cohorts are of predominantly European ancestry, so it is unknown whether the same sex-differential signals arise in other populations, and ancestrally diverse studies are needed for generalisability. Finally, the array interrogates only a fraction of the methylome and, as shown here, cannot support a valid interaction analysis of the X chromosome. Array methylation is a bulk, cell-averaged proportion that cannot separate the active from the inactivated X in females nor read the two alleles independently, so the differing male and female X compositions confound a bulk sex-interaction test. Allele-resolved, single-molecule approaches, in particular long-read sequencing, which reports CpG methylation on each read and can phase it to its parental haplotype and to the active or inactive X [43], together with variance models that accommodate X-inactivation, would allow the X chromosome to be analysed properly; broader assays such as EPIC v2 or whole-genome bisulfite sequencing would likewise extend coverage to regulatory regions the array omits. Region-level analysis (differentially methylated regions) was not pursued given the sparse, single-CpG nature of the signal, but may become informative at larger sample sizes. Greater power would also benefit the within-sex (Model 1) comparison, which is unbiased but, at present cohort sizes, the least able to reach significance.

Set against these limitations, the study has clear strengths: a large, multi-cohort and cross-platform sample analysed under a harmonised pipeline with bias-and-inflation correction and formal meta-analysis, and a deliberate contrast of stratified and interaction designs that clarifies why earlier main-effect analyses may have missed sex-specific signals. The broader implication is that sex should be modelled as a biological variable, and specifically as an interaction term in ALS methylation studies, rather than treated only as a covariate to be adjusted away.

## Conclusions

In a large, multi-cohort, interaction-based epigenome-wide association study of ALS, we identify seven autosomal sex-associated differentially methylated positions that are detectable by explicit case-by-sex interaction testing but not by sex-stratified comparison, and that are distinct from constitutive sex differences in blood methylation. Several of the implicated loci sit in pathways already central to ALS, namely neuronal excitability, energy metabolism and neuroinflammation, while others have no established connection to the disease; the effects are small throughout and await replication and functional validation. Beyond the specific loci, the work provides a sex-aware analytical reference for ALS methylation studies and argues for treating sex as a biological variable rather than a nuisance covariate.

## Ethics approval and consent to participate

All contributing studies were approved by the relevant institutional review boards or research ethics committees, and all participants provided written informed consent, in accordance with the Declaration of Helsinki. Project MinE was approved by the Trent Research Ethics Committee (08/H0405/60); the approving committees for the individual contributing cohorts are described in the supplementary cohort descriptions of our previous work [12] and in the SALSA-SGC cohort profile [22].

## Competing interests

A.A.-C. declares contracts with the MRC, NIHR and Darby Rimmer Foundation; consulting fees from Amylyx, Apellis, Biogen, Clene Therapeutics, Cytokinetics, GenieUs, GSK, Lilly, Mitsubishi Tanabe Pharma, Novartis, OrionPharma, Quralis, Sano, Sanofi and Vecalis, and is a judge for the Longitude Prize in ALS. A.A.K. declares contracts with the MRC (MR/Z505705/1), the Motor Neurone Disease Association (MNDA), National Institute for Health and Care Research (NIHR) Maudsley Biomedical Research Centre, Amyotrophic Lateral Sclerosis (ALS) Association Milton Safenowitz Research Fellowship, Darby Rimmer MND Foundation, LifeArc, the Dementia Consortium, and Black Swan Biotech; equipment by NIHR Maudsley Biomedical Research Centre; and consulting fees from the UK National Endowment for Science, Technology and the Arts (NESTA). L.H.v.d.B. reports grants from the Netherlands ALS Foundation, the Netherlands Organization for Health Research and Development (Vici scheme and the STRENGTH project under the EU Joint Programme – Neurodegenerative Disease Research, JPND) and the European Community’s Health Seventh Framework Programme (EuroMO-TOR, grant agreement 259867), and personal fees from Calico, Cytokinetics and Takeda and non-financial support from Orion and Orphazyme outside the submitted work. C.E.S. reports grants from AveXis, Eli Lilly, Chronos Therapeutics and Vertex Pharmaceuticals during the conduct of the study, and from QurAlis and Biogen outside the submitted work. The remaining authors declare no competing interests.

## Acknowledgements

We thank people with MND and their families for their participation, and the Project MinE ALS Sequencing Consortium and the contributing cohorts for data access. Samples used in this research were in part obtained from the UK National DNA Bank for MND Research, funded by the MND Association and the Wellcome Trust; we acknowledge sample management undertaken by Biobanking Solutions funded by the Medical Research Council at the Centre for Integrated Genomic Medical Research, University of Manchester. This work used the King’s College London CREATE high-performance computing facility. The authors acknowledge the National Institute for Health and Care Research (NIHR) Biomedical Research Centre at South London and Maudsley NHS Foundation Trust and King’s College London, and Health Data Research UK. The views expressed are those of the authors and not necessarily those of the NHS, the NIHR, King’s College London or the Department of Health and Social Care.

## Funding

Y.J.K. is funded by the Longitude Prize. A.A.K. is funded by the Longitude Prize, the MRC (MR/Z505705/1), the Motor Neurone Disease Association (MNDA), the National Institute for Health and Care Research (NIHR) Maudsley Biomedical Research Centre, the Amyotrophic Lateral Sclerosis (ALS) Association Milton Safenowitz Research Fellowship, the Darby Rimmer MND Foundation, LifeArc, the UK National Endowment for Science, Technology and the Arts (NESTA), and the Dementia Consortium. Equipment was funded by the NIHR Maudsley Biomedical Research Centre. This work was partly supported by an EU Joint Programme – Neurodegenerative Disease Research (JPND) project. The project is supported through the UK MND Research Institute and the following funding organisations under the aegis of JPND (www.jpnd.eu): United Kingdom, Medical Research Council (MR/L501529/1; MR/R024804/1) and Economic and Social Research Council (ES/L008238/1); and through the Motor Neurone Disease Association, My Name’5 Doddie Foundation, MND Scotland, LifeArc, the Alan Davidson Foundation, and the Darby Rimmer Foundation. This study represents independent research partly funded by the NIHR Biomedical Research Centre at South London and Maudsley NHS Foundation Trust and King’s College London, Rosetrees Trust, and the Medical Research Council, part of UK Research and Innovation.

## Data and code availability

The analysis code for the sex-stratified and interaction EWAS pipelines, together with the derived-result tables underlying each figure and table, is archived on Zenodo at https://doi.org/10.5281/zenodo.22689235. Full genome-wide summary statistics for both models are available from the corresponding author on reasonable request. Individual-level Project MinE methylation and phenotype data are available to researchers through the Project MinE consortium under its data-access procedures.

## Appendix Project MinE ALS Sequencing Consortium

Philip Van Damme, Philippe Corcia, Philippe Couratier, Patrick Vourc’h, Orla Hardiman, Russell L. McLaughlin, Marc Gotkine, Vivian Drory, Nicola Ticozzi, Vincenzo Silani, Jan H. Veldink, Leonard H. van den Berg, Mamede de Carvalho, Jesus S. Mora Pardina, Monica Povedano, Peter M. Andersen, Markus Weber, Nazli A. Başak, Ammar Al-Chalabi, Chris E. Shaw, Pamela J. Shaw, Karen E. Morrison, John E. Landers, Jonathan D. Glass and Clifton L. Dalgard.

## Supplementary tables

**Table S1:**
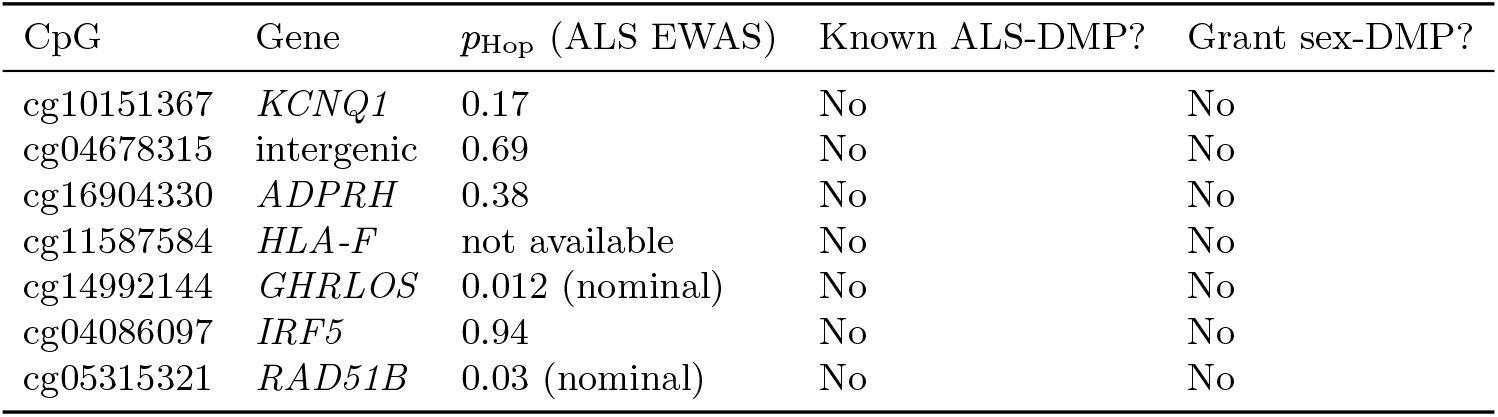
Novelty of the seven robust saDMPs relative to published catalogues: our previous blood ALS EWAS meta-analysis [12] (which reported 45 ALS-DMPs at 42 genes; overlap assessed against their full summary statistics for 724,712 CpGs) and the Grant et al. general-population autosomal sex-DMPs [17] (396 reported; the 395 available in the released list were tested here). *p*_Hop_ is the ALS main-effect meta-analysis *P* ; none of the seven reached genome-wide significance (*P <* 9 × 10^*−*8^) and two (*GHRLOS, RAD51B*) showed only nominal ALS association. None coincided with a general-population sex-DMP at CpG or gene level. cg11587584 (*HLA-F*) is absent from those summary statistics and so could not be compared. The seven positions are therefore detectable only by the ALS-by-sex interaction test, not by a conventional main-effect ALS EWAS or a sex-DMP scan.

| CpG | Gene | $p_{\text{Hop}}$ (ALS EWAS) | Known ALS-DMP? | Grant sex-DMP? |
| --- | --- | --- | --- | --- |
| cg10151367 | *KCNQ1* | 0.17 | No | No |
| cg04678315 | intergenic | 0.69 | No | No |
| cg16904330 | *ADPRH* | 0.38 | No | No |
| cg11587584 | *HLA-F* | not available | No | No |
| cg14992144 | *GHRLOS* | 0.012 (nominal) | No | No |
| cg04086097 | *IRF5* | 0.94 | No | No |
| cg05315321 | *RAD51B* | 0.03 (nominal) | No | No |

**Table S2:** Genomic inflation factor (*λ*) before and after *bacon* correction, by model and stratum, for the primary autosomal analysis; PCs is the tuned number of residual principal components selected for that fit (the X-chromosome analysis was tuned separately and is not shown here). Model 2 (ALS-by-sex interaction) is shown as *λ* per stratum with the inverse-variance meta-analysis at *λ* = 1.05; Model 1 (sex-stratified case/control) is shown as the sample-size-normalised *λ*_1000_ per stratum and sex. Residual inflation persists in the small AUS1 stratum but does not affect the conclusions, because Model 1 yielded no genome-wide saDMPs and the Model 2 meta-analysis is well calibrated.

| *Model 2: ALS × sex interaction ($\lambda$)* |  |  |  |  |
| --- | --- | --- | --- | --- |
| Stratum | $n$ | PCs | pre-*bacon* | post-*bacon* |
| MinE 450k | 4,354 | 10 | 1.06 | 1.02 |
| MinE EPIC | 3,505 | 10 | 1.07 | 1.02 |
| AUS1 | 1,083 | 30 | 1.44 | 1.03 |
| Meta (IVW) | 8,942 | – | – | 1.05 |
| *Model 1: sex-stratified case/control ($\lambda_{1000}$)* |  |  |  |  |
| Stratum (sex) | $n$ | PCs | pre-*bacon* | post-*bacon* |
| MinE 450k (M) | 2,541 | 30 | 1.16 | 1.02 |
| MinE 450k (F) | 1,813 | 25 | 1.15 | 1.02 |
| MinE EPIC (M) | 1,987 | 10 | 1.16 | 1.03 |
| MinE EPIC (F) | 1,518 | 30 | 1.24 | 1.03 |
| AUS1 (M) | 600 | 30 | 1.83 | 1.08 |
| AUS1 (F) | 483 | 30 | 3.44 | 1.20 |
| Sex-diff. meta | 8,942 | – | – | 1.00 |

## Supplementary figures

**Figure S1:**
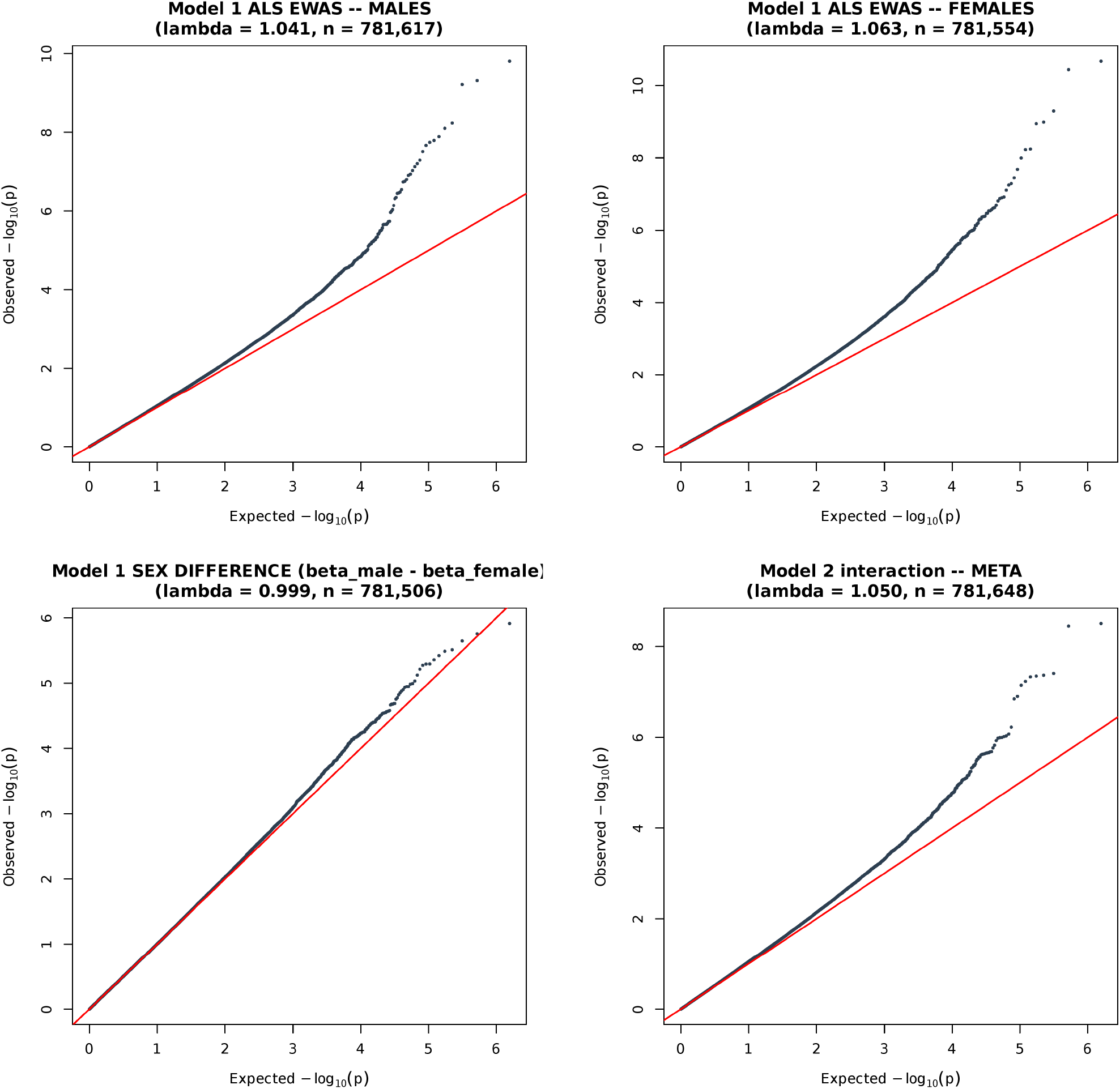
Quantile–quantile plots of the meta-analysed association *P* values after PC tuning and *bacon* correction; in every panel the observed − log_10_ *P* is plotted against the quantiles expected under the null, and *λ* is the genomic inflation factor of those *P* values. For Model 1 the three panels are the two sex-stratified case/control EWAS that the model fits separately, (A) males (*λ* = 1.04) and (B) females (*λ* = 1.06), and (C) the between-sex difference test that contrasts them (*λ* = 1.00); showing all three indicates that the difference test is well calibrated because each underlying per-sex EWAS is, rather than because two inflated distributions cancel. Panel (D) is the Model 2 interaction meta-analysis (*λ* = 1.05). Note that (C) plots the *P* values of the formal test of *β*_male_ = *β*_female_, which is uniform under the null because the two estimates come from disjoint samples; it is not a plot of the effect-size difference itself.

**Figure S2:**
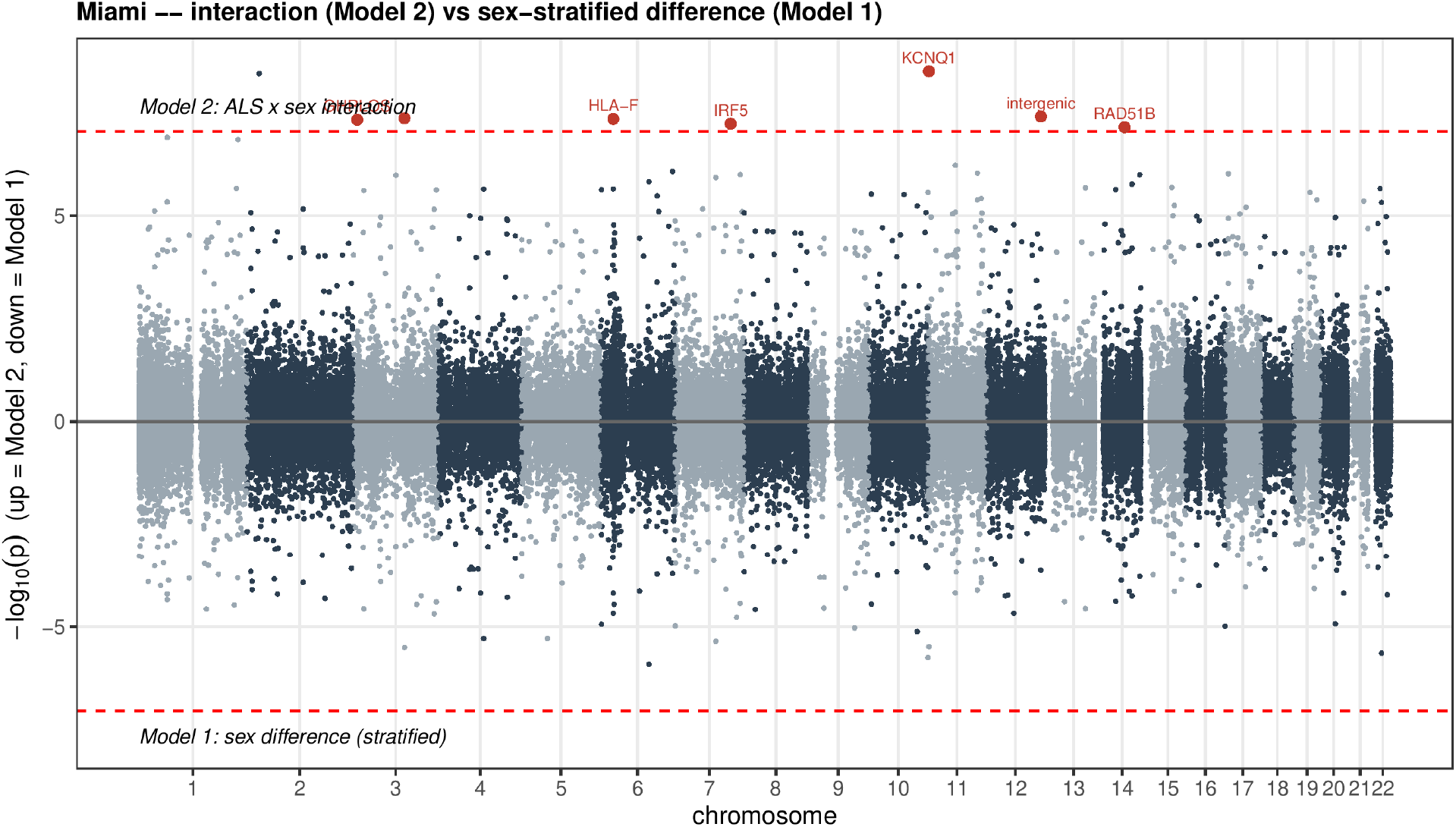
Miami plot contrasting the sex-stratified difference test (Model 1, plotted downward) with the ALS-by-sex interaction meta-analysis (Model 2, plotted upward) on a shared genomic axis. The interaction model recovers genome-wide signals that the stratified difference test does not, illustrating the central methodological point of the study.

**Figure S3:**
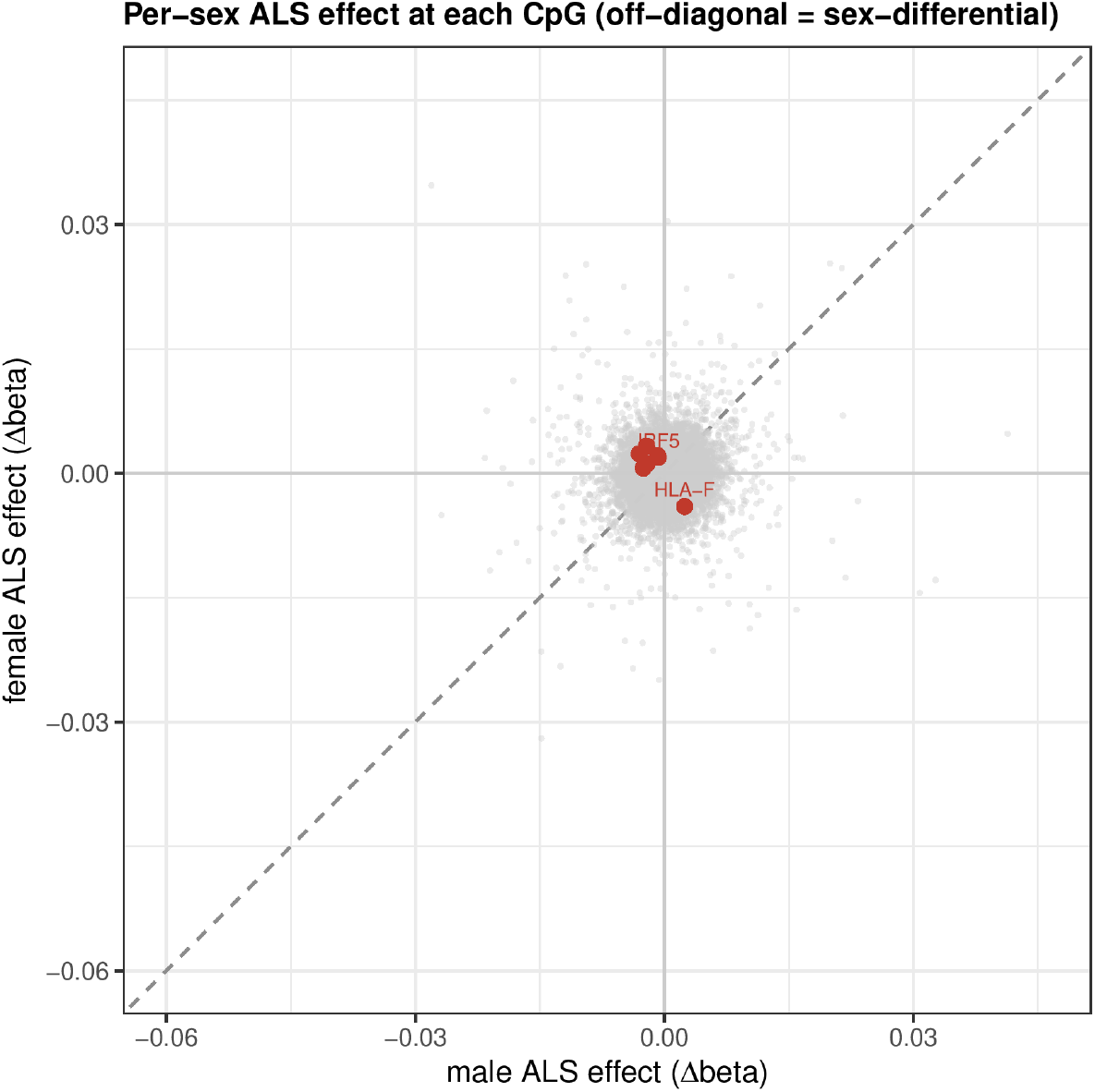
Per-sex ALS case/control effect at each autosomal CpG (male on the *x*-axis, female on the *y*-axis; grey points subsampled). CpGs on the dashed identity line change concordantly in both sexes, whereas the seven robust saDMPs (red) lie off the diagonal, the signature of a sex-differential effect.

## Notes

https://doi.org/10.5281/zenodo.22689235

